# To hum or not to hum: Neural transcriptome signature of courtship vocalization in a teleost fish

**DOI:** 10.1101/2020.10.05.327304

**Authors:** Joel A. Tripp, Ni Y. Feng, Andrew H. Bass

**Affiliations:** Department of Neurobiology and Behavior, Cornell University, Ithaca, NY; Department of Integrative Biology, University of Texas-Austin, Austin, TX; Yale University School of Medicine, New Haven, CT

**Keywords:** vocalization, motoneuron, preoptic area, transcriptome, gene expression, circadian, neuropeptide, synaptic transmission, ion channel, metabolism

## Abstract

For many animal species, vocal communication is a critical social behavior, often a necessary component of reproductive success. In addition to the role of vocal behavior in social interactions, vocalizations are often demanding motor acts. Through understanding the genes involved in regulating and permitting vertebrate vocalization, we can better understand the mechanisms regulating vocal and, more broadly, motor behaviors. Here, we use RNA-sequencing to investigate neural gene expression underlying the performance of an extreme vocal behavior, the courtship hum of the plainfin midshipman fish (*Porichthys notatus*). Single hums can last up to two hours and may be repeated throughout an evening of courtship activity. We asked whether vocal behavioral states are associated with specific gene expression signatures in key brain regions that regulate vocalization by comparing transcript levels in humming versus non-humming males. We find that the circadian-related genes *period3* and *Clock* are significantly upregulated in the vocal motor nucleus and preoptic area-anterior hypothalamus, respectively, in humming compared to non-humming males, indicating that internal circadian clocks may differ between these divergent behavioral states. In addition, we identify suites of differentially expressed genes related to synaptic transmission, ion channels and transport, hormone signaling, and metabolism and antioxidant activity that may permit or support humming behavior. These results underscore the importance of the known circadian control of midshipman humming and provide testable candidate genes for future studies of the neuroendocrine and motor control of energetically demanding courtship behaviors in midshipman fish and other vertebrate groups.

## 1. Introduction

Communication is a critical aspect of social behavior, and for many species, vocal communication is a necessary step in finding or attracting mates^1,2^. Vocalization is controlled by a hierarchy of systems that includes environmental context, intrinsic drive, central pattern generators, and peripheral sound-producing organs^3^. In the case of advertisement vocalization, factors influencing the upper levels of this hierarchy (context and drive) may be related to the likelihood that an appropriate audience is present (e.g. correct season and time of day to attract a mate) and an individual’s own reproductive status signaled by hormonal state. At the most basic level of behavioral action, these vocalizations often require complex or demanding motor control^4–6^. To better understand both the neuroendocrine and motor mechanisms driving vocal communication, we have studied the plainfin midshipman (*Porichthys notatus*), a teleost fish capable of producing advertisement calls that are extreme in terms of duration and spectro-temporal simplicity^7,8^. The neural control of vocalization in this species is also well-established^4,9^. Male midshipman develop into one of two alternate morphs: type I males that aggressively defend nests, vocally court females, and provide parental care or type II males that do not engage in these behaviors, and instead reproduce by stealing fertilizations at the nests of type I males^10^. Here, we focus on type I males, and the neural network controlling vocalization. Courting type I males use hums to attract females to their nest to spawn^1^. These calls are relatively simple in fine structure (multi-harmonic, sinusoidal-like waveform), but may last up to two hours in duration and can be produced repetitively during a single night of courtship^8^. Courtship humming is highly seasonal and under circadian control^8^, occurring at night during the late spring and summer breeding season^7,10,11^. Males chorus at night, with hums dominating the underwater soundscape for much of an evening in the habitat where midshipman nest during the breeding season^7^.

Type I males have an expansive vocal-motor system^12,13^ in comparison to type II males and females that are only known to produce brief, agonistic calls. This includes enlarged muscles attached to the walls of the swim bladder, whose vibrations produce sound pulses that are directly controlled by a well-characterized hindbrain vocal pattern generating circuit^9^. The final node in this circuit is the vocal motor nucleus (VMN), a paired midline nucleus. Each VMN contains close to 2000 motor neurons that innervate the ipsilateral vocal muscle attached to the swim bladder wall (Figure 1A). The paired VMN display extraordinary synchrony in their firing, with population activity matching 1:1 the firing of individual neurons, the simultaneous contraction of the paired vocal muscles, and the onset of single sound pulses^4,14^ (∼100 Hz at 16°C). Previously, we identified changes in the transcriptional profile of the VMN over both seasonal and daily time scales^5^ and differential transcript expression during spawning across male morphs and tactics in the preoptic area-anterior hypothalamus (POA-AH)^15^.

**FIGURE 1.**
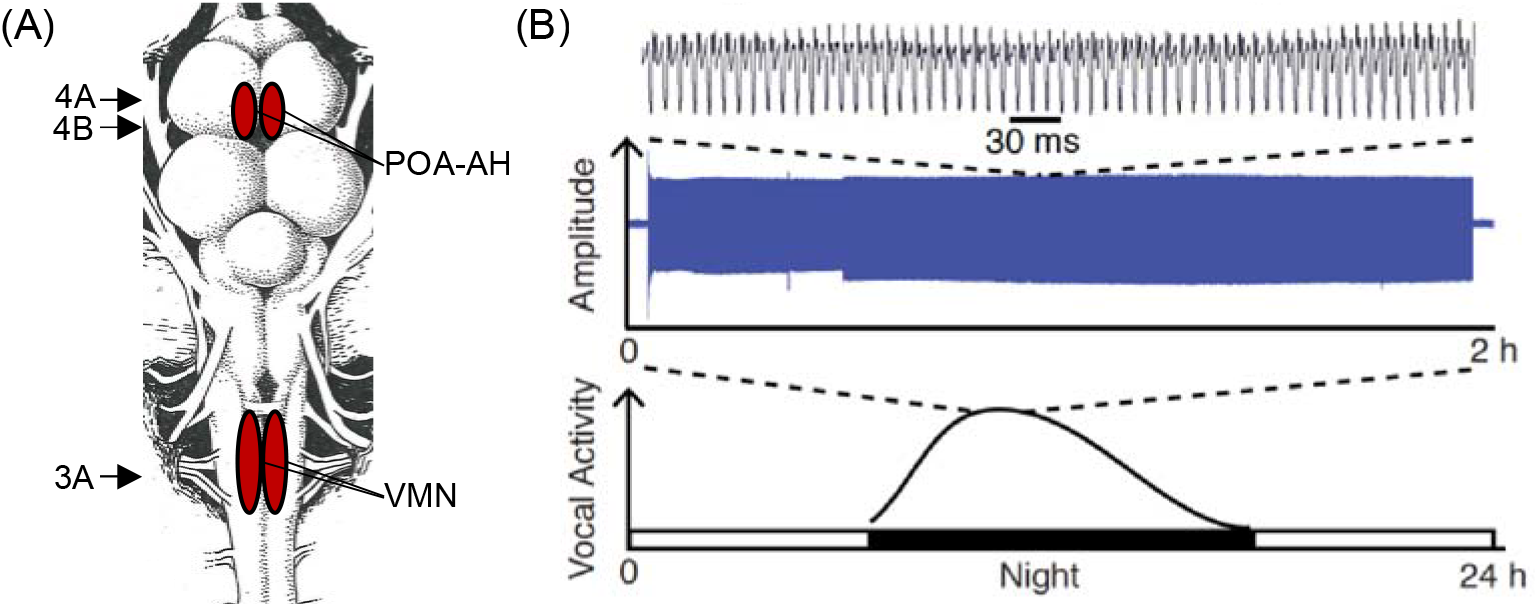
Midshipman vocal system. (A) Overhead view of midshipman brain. Arrows indicate levels of cresyl-stained tissue sections shown in Figures 3 and 4. (B) Recording of midshipman courtship hum shown at multiple time scales (from Feng and Bass, 2016). Hums are produced nearly exclusively at night during the summer breeding season. A single hum can last up to two hours in duration, and multiple hums are produced repetitively through the course of a night.

Interestingly, while all type I males have the neuromuscular capability to hum (e.g., see Bass et al., 1994; Brantley and Bass, 1994), only some are observed to hum regularly in captivity under semi-natural conditions, while others never hum (e.g., see Feng & Bass 2016 and Supplemental Information therein). These divergent behavioral states are comparable to observations of vocal activity in the midshipman’s natural habitat (see McIver et al., 2014); during some evenings of monitoring nesting sites, some type I males do not hum and only emit agonistic grunts and growls (see McIver et al., 2014; M. Marchaterre and A. Bass, unpublished observations). Taking advantage of these divergent behavioral states in captive populations, we wanted to know whether the behavioral state of humming is paralleled by gene expression differences in brain regions controlling neuroendocrine and motor aspects of humming behavior. First, we hypothesized that the VMN of humming males may be “primed” with a suite of genes whose function supports the neurophysiological and energetic demands of producing the highly synchronous and long-duration motor command for humming. Second, we hypothesized that because the POA-AH is a major vertebrate neuroendocrine center (Figure 1A) whose neurons have been shown to modulate midshipman vocalization^16,17^, genes differentially expressed in the POA-AH of humming males likely play a role in initiating or permitting the humming behavioral state.

We used RNA-sequencing (RNAseq) to identify transcriptional changes that occur during humming in both the VMN and POA-AH of type I male midshipman. We found that the vocally active behavioral state of humming is characterized by differential expression of a suite of functionally important genes supporting synaptic transmission, ion channels and transport, hormone signaling, and metabolism and antioxidant activity—categories that have been the focus of earlier studies of vocalization in midshipman or have been associated with courtship vocalization^5,15,18,19^. We also found that the circadian genes *period3* (*per3*) and *circadian locomotor output cycles kaput-like* (*Clock*) were differentially expressed between humming and non-humming males in the VMN and POA-AH, respectively; an unexpected but exciting result given that midshipman vocalization is under circadian control^8^. Together, these findings underscore the importance of circadian control of vocal communication behaviors critical for reproduction in this and other vertebrate species^8^, and identify a list of candidate genes that may play a role in promoting, permitting, or patterning an extreme vocal display.

## 2. Materials and Methods

### 2.1 Animal subjects

Type I male midshipman were collected from nests in Washington state in June 2013 and housed at the University of Washington Big Beef Creek field station. All procedures were approved by the Institutional Animal Care and Use Committees of the University of Washington and Cornell University.

### 2.2 Tissue collection

Following behavioral methods previously used to promote courtship humming in captivity^15,20,21^, midshipman type I males were housed in large (2m x 2m x 1m) outdoor tanks in ambient light and temperature. Each tank contained six or eight artificial nests (one for each fish) made of a ceramic plate resting on bricks (Figure 2A, B). Vocal behavior was monitored using custom-built hydrophones (Bioacoustics Research Program, Cornell Laboratory of Ornithology) connected to digital audio recorders (Olympus LS-12). Humming males (n=4, 50-120 g body weight, 15.9-20.3 cm standard length) were collected at least 30 minutes after the onset of humming (Figure 2C). Experimenters confirmed that a male had been humming before sacrifice on the basis of swim bladder inflation^22^. Humming males are easily identified, as they are buoyant and float to the surface of the water when artificial nest tops are removed. Non-humming males (n=4, 90-105 g body weight, 18.5-20.3 cm standard length) were collected from the same aquaria that housed humming males at similar time points (time of sacrifice 10:15pm-12:35am for humming and 10:35pm-1:18am for non-humming).

**FIGURE 2.**
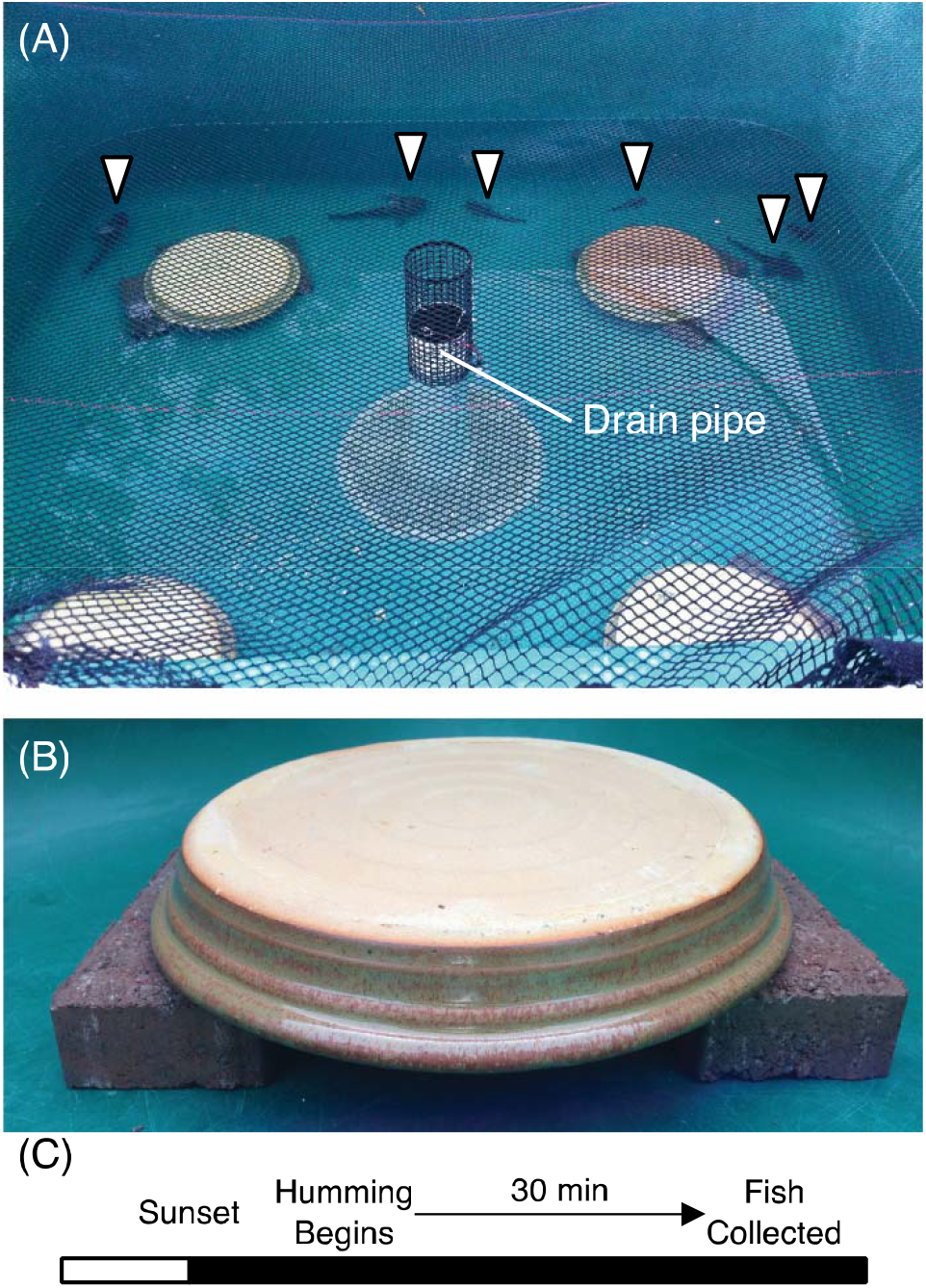
Experiment overview. (A) Example of outdoor tank used to house fish. Arrows indicate fish swimming outside of nests (tan-colored ceramic plate resting on bricks). (B). Close-up image of artificial nest. Photos taken during the day to ensure visibility. (C) Time course of animal collection. Courtship humming began after sunset. Humming individuals were identified and collected at least 30 min after onset of vocalization. Non-humming males were collected at similar time points.

To collect brain tissue, fish were first deeply anesthetized in 0.025% benzocaine (Sigma Aldrich, St. Louis, MO), and then exsanguinated through the heart. Brains were quickly removed, transected at the midbrain-hindbrain boundary, stored in RNAlater (Life Technologies, Carlsbad, CA) at 4° C overnight, and then at −20° C. Tissues were shipped overnight on dry ice to Cornell University (Ithaca, NY) and stored at −20 or −80° C until use.

### 2.3 RNA extraction and sequencing

Brains in RNAlater were thawed on ice, then the VMN and POA-AH were dissected in RNAlater as previously described^5,15,23^. Briefly, to collect the VMN (Figure 3A), fine forceps were used to separate the entire nucleus from surrounding hindbrain under a dissecting microscope. To collect the POA-AH (Figure 4A, B), the telencephalon was removed, and the POA-AH was cut away from the remaining brain under a dissecting microscope. RNA was extracted in Trizol (Invitrogen, Carlsbad, CA), treated with DNase I (Invitrogen), and reverse transcribed using Superscript III (Invitrogen) following manufacturer’s protocols. 3.22-9.8μg of cDNA per tissue sample was sent to Polar Genomics (Ithaca, NY) for strand-specific library preparation. Barcoded libraries were made for each tissue type from each individual. Sequencing was performed using the Illumina NextSeq 500 by the Cornell University Biotechnology Resource Center Genomics Facility. Reads were deposited in the NCBI Sequence Read Archive under BioProject Accession Number PRJNA665270^24^.

**FIGURE 3.**
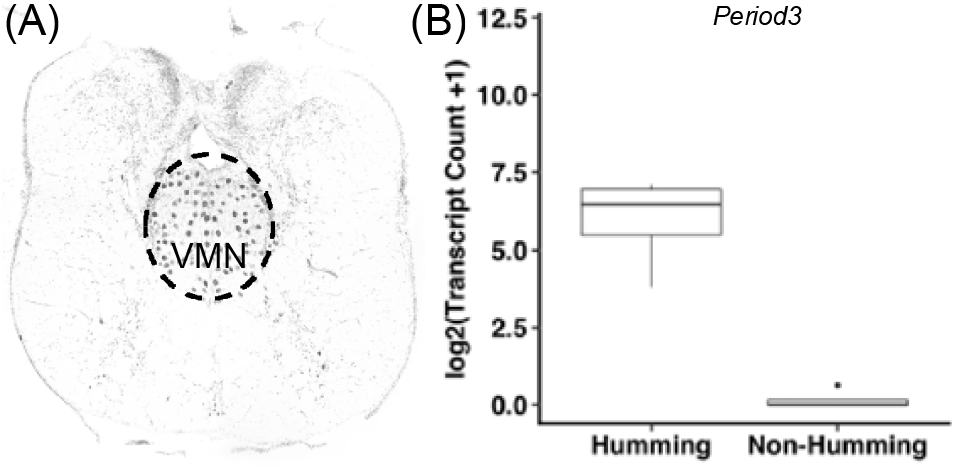
*Period3* expression in humming and non-humming male midshipman VMN. (A) Coronal section through hindbrain at the level of the paired VMN contiguous along the midline shown in Figure 1A. (B) Boxplot shows log_2_ of *per3* transcript counts+1 in humming and non-humming males (N=4 for each group). Dark lines within boxes indicate median values, lower and upper box edges indicate first and third quartiles, and whiskers extend to minimum and maximum values. Dot indicates outlier value (more than 1.5 times the interquartile range from the third quartile).

**FIGURE 4.**
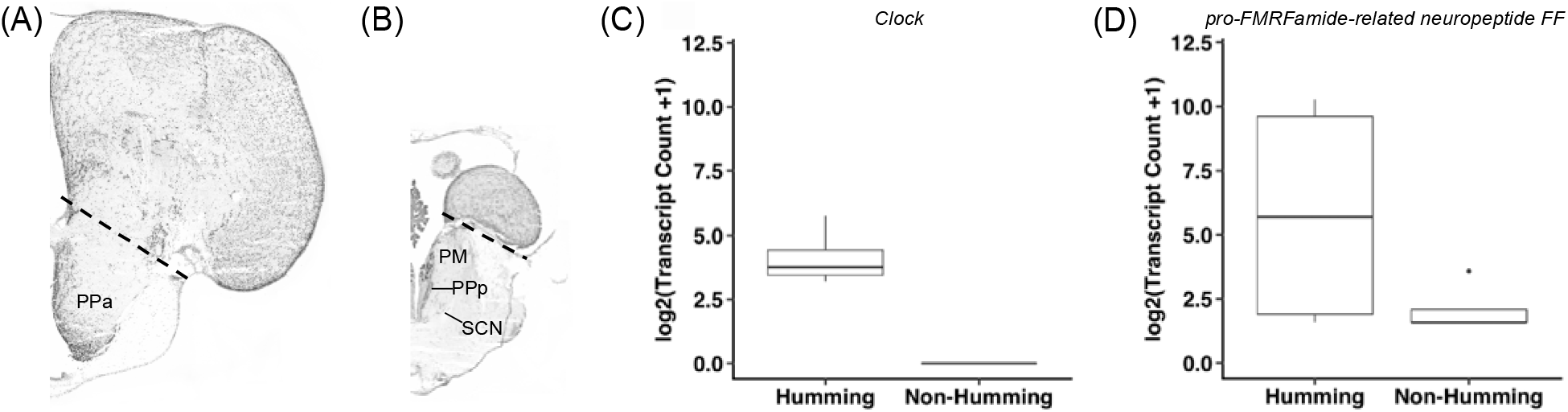
Transcript expression in humming and non-humming male midshipman POA-AH. (A) Coronal sections through forebrain at the level of the anterior (A) and posterior (B) POA-AH indicated in Figure 1A. Dashed lines indicate tissue cut to collect POA-AH for RNA-sequencing. Abbreviations: PM, magnocellular preoptic nucleus; PPa, anterior parvocellular preoptic nucleus; PPp, posterior parvocellular preoptic nucleus; SCN, suprachiasmatic nucleus. Boxplots of (C) *Clock* and (D) Neuropeptide FF expression in humming and non-humming males (N=4 for each group). Boxplots show log_2_ of transcript counts+1. Dark lines within boxes indicate median values, lower and upper box edges indicate first and third quartiles, and whiskers extend to minimum and maximum values. Dot indicates outlier value (more than 1.5 times the interquartile range from the third quartile).

### 2.4 Reference transcriptome

The methods followed here were adopted from prior studies of the midshipman VMN^5^ and POA-AH^15^. Following sequencing, Illumina quality filtering removed read pairs in which either read was poor quality. Adapter sequences and low quality nucleotides were removed from reads using Trimmomatic (v0.32)^25^. POA-AH and VMN reads from the humming individual with the most total reads were used to assemble a reference transcriptome using Trinity (v2.1.1)^26^. A humming individual was chosen to ensure the assembly of potential rare transcripts that may only be expressed during humming. The reference assembly was assessed and filtered using Transrate (v0.32)^27^ and completeness was assessed using the BUSCO (v3.0.2) Actinopterygii dataset^28^. To annotate the reference assembly, Transdecoder (v2.1.0)^29^ was used to identify open reading frames and generate peptide sequences which were submitted to the blastp refseq_protein database with an e-value cutoff of 1 × 10^−10^, then annotated using Blast2Go^30^. Additional functional annotation information of differentially expressed transcripts was obtained by manual search for the identified protein function in the UniProtKB database (uniprot.org). Sequences that lacked a vertebrate BLAST hit were removed.

### 2.5 Statistical analysis

Differential expression analyses were conducted using edgeR (Bioconductor, v3.16.5)^31,32^. Kallisto^33^ was used to estimate transcript abundance, abundance estimates were imported using tximport (Bioconductor, v1.2.0)^34^, and the data were fit to a generalized linear model. Comparisons were made between humming and non-humming males for POA-AH and VMN tissue libraries. False discovery rate (FDR)<0.05 was used as a cut-off for significance^35^. Differences between groups were reported as log2-fold change (logFC) in transcript expression. Analysis of Gene Ontology (GO) term enrichment was conducted using Blast2Go^30^. Fisher’s exact test was used to identify GO terms significantly enriched among genes found to be differentially expressed in either VMN or POA-AH with an FDR cutoff of 0.1.

## 3. Results

### 3.1 Reference transcriptome

Trinity assembly resulted in a transcriptome containing 406,689 sequences with a mean length of 642.06bp, Transrate score of 0.07088, with 77.7% complete, 8.9% fragmented, and 13.4% missing BUSCOs. Filtering with Transrate maintained 174,421 sequences with a mean length of 919.72bp, Transrate score of 0.28012, with 74.6% complete, 8.8% fragmented, and 16.6% missing BUSCOs. The final reference transcriptome contained 53,612 sequences each with identified open reading frame and at least one vertebrate BLAST hit.

### 3.2 Differential expression—VMN

We found that 177 transcripts were significantly differentially expressed (FDR<0.05) between humming and non-humming males in the VMN (Supplementary Table 1). No GO terms were found to be significantly enriched at the FDR<0.1 level. Among the differentially expressed transcripts was the circadian-related gene *per3*, which was upregulated in humming males (Figure 3). In addition, we identified twelve transcripts involved in synaptic transmission, eight related to ion channels and transport, and four involved in metabolism and antioxidant activity that were differentially expressed between humming and non-humming animals (Table 1).

**TABLE 1.** Differentially expressed transcripts in VMN

| *Category* | *Blast2Go Description* | *Expression in Hummers*<br>*(logFC, FDR)* |
| --- | --- | --- |
| Circadian Rhythm | *period circadian homolog 3 isoform X1* | Up (8.45, 0.00012) |
| Synaptic<br>Transmission | *AP-2 complex subunit mu isoform X2* | Up (8.27, 7.77*10<sup>-20</sup>) |
|  | *C-Jun-amino-terminal kinase-interacting 2 isoform X2* | Up 13.19 (1.61*10<sup>-8</sup>) |
|  | *ELKS Rab6-interacting CAST family member 1 isoform X2* | Up (2.85, 0.00016) |
|  | *gamma-aminobutyric acid receptor subunit beta-2 isoform X2* | Up (2.11, 0.00083) |
|  | *leucine-rich repeat-containing 4B-like* | Up (7.40, 0.0012) |
|  | *metabotropic glutamate receptor 3* | Up (6.28, 0.0076) |
|  | *vacuolar sorting-associated 33A* | Down (7.98, 0.010) |
|  | *synapsin-2* | Down (5.60, 0.017) |
|  | *syntaxin-16 isoform X2* | Up (8.45, 0.00074) |
|  | *syntaxin-binding 5-like isoform X6* | Down (8.04, 0.00050) |
|  | *syntaxin-binding 5-like isoform X10* | Up (5.14, 0.039) |
|  | *vacuolar sorting-associated 33A* | Down (7.98, 0.010) |
| Ion Channels &<br>Transport | *inactive dipeptidyl peptidase 10 isoform X1* | Down (2.58, 0.022) |
|  | *probable voltage-dependent N-type calcium channel subunit alpha- partial* | Up (6.05, 0.047) |
|  | *solute carrier family 22 member 15 isoform X1* | Down (6.02, 0.0023) |
|  | *transmembrane and coiled-coil domain-containing 3* | Up (11.78, 0.0091) |
|  | *unc-80 homolog* | Up (4.18, 0.00077) |
|  | *voltage-dependent calcium channel gamma-4 subunit-like* | Up (4.42, 0.0015) |
|  | *voltage-dependent calcium channel subunit alpha-2 delta-1* | Down (5.87, 0.036) |
|  | *voltage-dependent calcium channel gamma-2 subunit* | Up (7.76, 0.049) |
| Metabolism &<br>Antioxidant Activity | *apolipo C-II-like* | Up (6.73, 0.016) |
|  | *fructose-2,6-bisphosphatase TIGAR B-like isoform X1* | Up (2.91, 0.0085) |
|  | *lipid phosphate phosphatase-related type 5-like isoform X2* | Down (5.12, 0.036) |
|  | *L-lactate dehydrogenase A chain* | Up (9.61, 0.0013) |

### 3.3 Differential expression—POA-AH

Seventy-one transcripts were significantly differentially expressed (FDR<0.05) between humming and non-humming males in the POA-AH (Supplementary Table 2). No GO terms were significantly enriched at the FDR<0.1 level. Differentially expressed transcripts included *Clock*, a gene involved in circadian rhythm, that was upregulated in humming males (Figure 4A). Additionally, we identified one transcript related to hormone signaling, *pro-FMRFamide-related neuropeptide FF* (Figure 4B), four involved in synaptic transmission, and five encoding ion channels or transporters that were differentially expressed between humming and non-humming animals (Table 2).

**TABLE 2.**
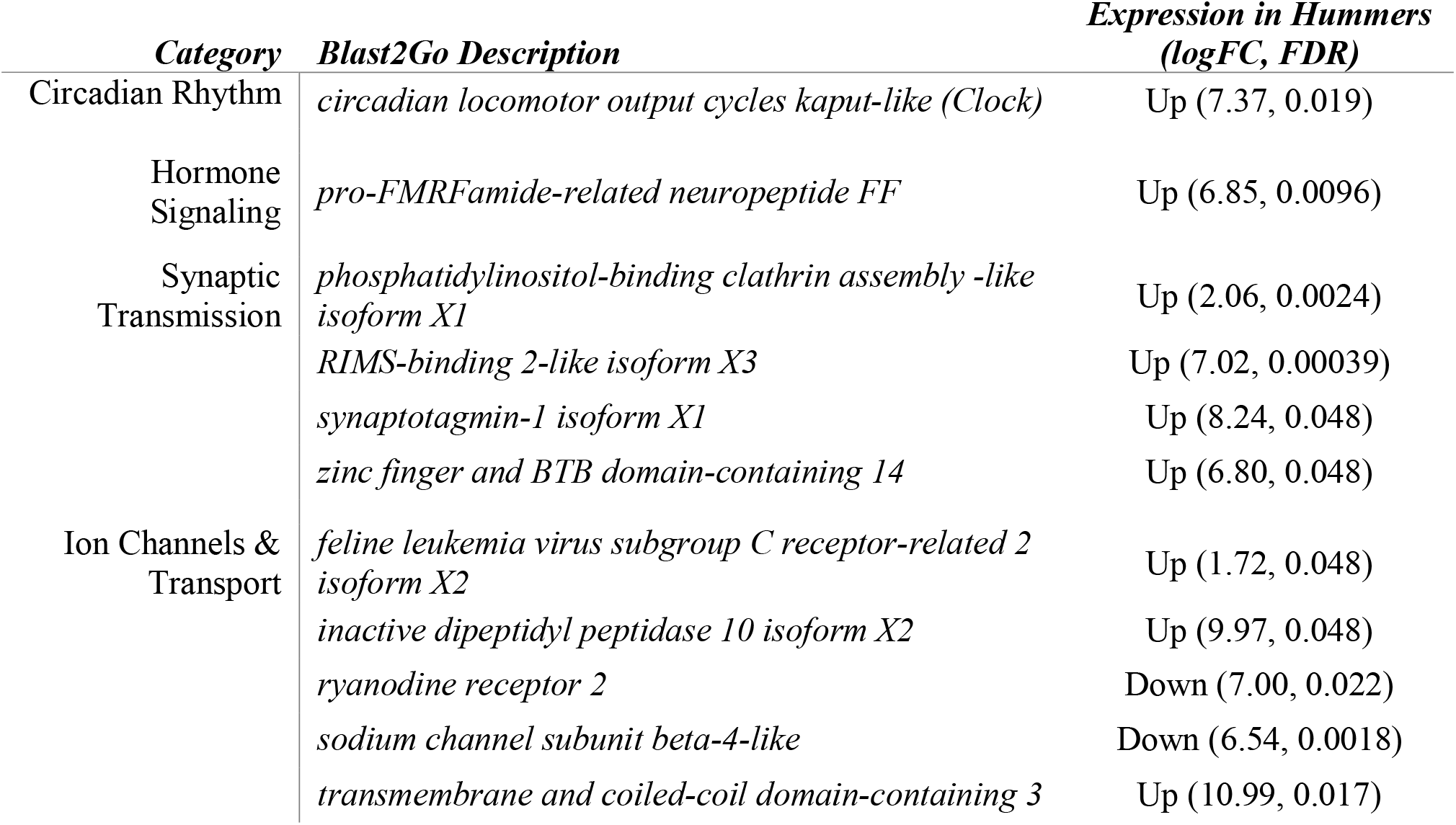
Differentially expressed transcripts in POA-AH

## 4. Discussion

Vocalization is a widely used signaling modality for vertebrate animal communication^2^. For males of many species, the ability to perform courtship songs and vocalizations are essential for reproductive success^2^. In this study, we investigated changes in brain gene expression within the neural network controlling courtship vocalization. The courtship hum of the midshipman fish results from swim bladder vibrations caused by paired vocal muscles that “generate more muscle contractions in an hour than any other known muscle”^36^. We chose to focus on two regions previously studied for their role in vocalization and other social behaviors: the VMN, the motor nucleus which directly innervates midshipman vocal muscle^37^, and the POA-AH, a neuroendocrine center found across vertebrate lineages which is involved in mating and produces neuropeptides which modulate midshipman vocalization^16,38–40^. Within each of these regions, we identified sets of differentially expressed transcripts with functions related to circadian rhythm, synaptic transmission, and ion channels and transport. Additionally, four differentially expressed transcripts with roles in metabolism and antioxidant activity were identified within the VMN, and one transcript related to hormone signaling in the POA-AH.

### 4.1 Relation to prior midshipman studies of the VMN

This study builds on our previous investigations of gene expression changes related to midshipman vocalization and mating behavior. The first examined gene expression across daily and seasonal timescales relevant to midshipman humming behavior and compared the VMN to the surrounding hindbrain^5^. That study identified several gene categories upregulated in the VMN compared to surrounding hindbrain tissue, including genes related to neurotransmission, steroid biosynthesis, neuropeptides/peptide hormones, thyroid hormone, neuromodulators, and antioxidants. Similarly, several genes related to neurotransmission, steroid receptors and metabolic enzymes, and peptide hormones and receptors had peak expression levels in the summer breeding season, or more specifically at summer night when humming occurs.

Feng et al. (2015) identified several categories of gene expression in the VMN that we further investigated here. However, somewhat surprisingly, we did not find overlap in the precise transcripts that were differentially expressed across tissue types and timescales of the previous study and across humming and non-humming individuals that are the focus here. It is important to emphasize key methodological differences between the two studies. The prior one made comparisons between hindbrain regions and across daily and seasonal timescales, while here we make comparisons within the VMN across behavioral states, namely humming and non-humming individuals collected at similar times in a natural light cycle. These key differences may explain the differences in our results. While the transcripts upregulated in VMN during the summer night, in general, are likely to play an important role in preparing the nucleus for the demands of humming behavior, our results would suggest that they do not further change expression once humming begins, but instead other additional, related transcripts may play key functional roles during humming behavior. Furthermore, we chose to collect tissue after only 30 min of humming to avoid activity induced gene expression^41^ to isolate the transcripts that best determine whether or not a fish would initiate humming. The short duration of humming likely explains the relatively few differentially expressed genes we observed.

Despite differences between our study and the prior one of the VMN, the general trends remain the same. For example, the previous study found several voltage-gated calcium channel subunits upregulated in VMN compared to surrounding hindbrain; here, we also found the transcripts *voltage-dependent calcium channel gamma-4 subunit-like, probable voltage-dependent N-type calcium channel subunit alpha-partial*, and *voltage-dependent calcium channel gamma-2 subunit* upregulated in humming males, though one subunit type, *voltage-dependent calcium channel subunit alpha-2 delta-1*, was downregulated in humming compared to non-humming males (Table 1). Further, the inhibitory transmitter gamma-aminobutyric acid (GABA) has been shown to play an important role in maintaining the extraordinary synchrony displayed by VMN neurons^4^. *GABA receptor subunit pi-like* was previously shown to peak in expression during the summer nights. Here, we found *gamma-aminobutyric acid* (GABA) *receptor subunit beta-2 isoform X2* to be significantly upregulated in humming males. Humming males also showed significant increases in *unc-80* expression. Unc-80 is part of the NACLN leak sodium channel that contributes to resting potential levels^42^. Increased NACLN expression could figure prominently into increasing the low excitability of VMN motor neuron at rest^4^ during long duration hums.

### 4.2 Relation to prior midshipman studies of the POA-AH

In another RNAseq study, we examined transcriptomic changes in the POA-AH related to spawning in male midshipman^15^. In that study, courting males were collected after they were found to be spawning, but because successful courtship requires a male to attract a female by humming, it was not possible to fully disentangle the gene expression changes related to spawning *per se* from those associated with related behaviors, including courtship humming along with other behaviors that occur throughout a spawning bout (e.g., aggression toward male intruders). The current focus on comparing humming males to non-humming males more specifically makes it possible to better identify gene expression changes related to courtship vocalization versus other spawning behaviors.

Only a single transcript related to hormone signaling, *pro-FMRFamide-related neuropeptide FF* was differentially expressed between humming and non-humming midshipman males. This result was surprising because the POA-AH has dense expression of neuropeptides and hormone receptors—including arginine-vasotocin (AVT) and isotocin (teleost homologues of mammalian arginine-vasopressin and oxytocin, respectively) and their receptors, galanin, stress-related peptides (corticotropin-releasing hormone and urocortin-3), somatotropin, androgen and estrogen receptors, the enzyme aromatase which converts testosterone to estradiol, as well as a receptor for the hormone melatonin, which promotes vocalization^8,15,18,19,39,40,43–46^. Further, hormone signaling was the single significantly enriched GO term in our study of the POA-AH during spawning^15^. Despite this, the only hormone signaling-related transcript that was differentially expressed in the present study, neuropeptide FF, is an interesting candidate for a role in vocal regulation.

Neuropeptide FF is found in the hypothalamus of rats, including the supraoptic and paraventricular nuclei^47^, considered to be at least partial homologues of the parvocellular and magnocellular populations of the teleost POA-AH^48^. Functionally, neuropeptide FF plays a role in pain perception, spatial behavior, and a wide range of physiological functions, including response to osmotic stress and regulation of blood pressure^47^. Intriguingly, intracerebroventricular injection of neuropeptide FF inhibits release of arginine-vasopressin in response to injection of hypertonic saline in mice^49^. As AVT inhibits type I male midshipman vocal-motor output (termed fictive calling) in *in situ* neurophysiological studies^16,39^, it is possible that neuropeptide FF plays a similar role here, reducing the release of AVT to prevent the inhibition of vocal output.

Neuropeptide FF also interacts with opioid signaling in the central nervous system. This is of note, as opioid antagonists inhibit fictive calling in an *in situ* neurophysiological preparation when pressure injected into the region of the midshipman’s hindbrain pattern generating circuit^50^. When administered intrathecally, neuropeptide FF has analgesic effects and potentiates opioid action, while intracerebroventricular injection results in anti-opioid effects and chronic infusion of neuropeptide FF into the ventricles of rats results in downregulation of the mu-opioid receptor^51^. These opposing studies seem to indicate that neuropeptide FF interactions with opioid signaling differ between the spinal cord and forebrain. The midshipman VMN sits on the hindbrain-spinal cord boundary, so it may be that neuropeptide FF is released from POA-AH neurons and acts locally or in the vocal pattern generating nuclei (VMN along with pacemaker and pre-pacemaker nuclei) with faciliatory action on the opioid system, similar to intrathecal administration in mice.

While neuropeptide FF expression has not yet been described in the teleost POA-AH, significant neuropeptide FF-like immunoreactivity has been described in the preoptic regions and hypothalamus of lamprey (*Petromyzon marinus*); anuran, urodele, and gymnophionan amphibians; and geckos (*Gekko gecko*)^52–55^. Dense fibers were also observed in the hypothalamus, hindbrain and spinal cord of these species, consistent with possible interactions with AVT and opioid signaling at those levels. Whether neuropeptide FF enhances or inhibits opioid signaling in midshipman is unclear, however, it’s interactions with neuromodulatory systems known to play important roles in midshipman vocal behavior make it an intriguing candidate for future study.

As in our previous study of the POA-AH during spawning^15^, we did not find differential expression of either AVT or isotocin. This was surprising as the POA-AH is the primary source of these peptides in the midshipman brain^39,40^, and they influence midshipman vocal-motor physiology in a male morph-specific manner^16^. Specifically, AVT reduces evoked vocal-motor output in type I males in an *in situ* neurophysiology preparation (isotocin has a similar effect in type II males and females). One possible explanation is that these peptides influence only specific types of vocalization. While we focused on courtship humming here, the fictive vocalizations evoked during *in vivo* physiology studies typically resemble the much shorter grunts and growls produced in agonistic contexts^7,10^. It may be that the nonapeptides exert their morph-specific effects on these agonistic signals, but do not play a role in courtship vocalization. Alternatively, it may be that AVT is produced at similar levels in humming and non-humming males, but its release is blocked in humming animals (see above), disinhibiting vocalization. Another explanation could be that changes in transcript levels of these peptides require longer than 30 min of humming behavior.

### 4.3 Role of circadian genes in vocalization

Our results also highlight the importance of circadian control of vocal behavior. Midshipman humming, like the vocal behavior of many bird species^56–58^, is under circadian control^8^. Here, we found two genes involved in circadian time keeping were differentially expressed across humming and non-humming midshipman. *Per3* was upregulated in the VMN of humming males, while *Clock* was significantly upregulated in the POA-AH of humming males.

Circadian time keeping genes are highly conserved among both vertebrates and invertebrates, acting through transcription feedback loops^59^. In vertebrates, *Clock* forms a heterodimer with *BMAL1* to activate transcription of Cryptochrome and Period genes, which in turn repress expression of *Clock* and *BMAL1*. Period genes peak in expression six hours after light onset in the suprachiasmatic nucleus of mice, which is the main mammalian time-keeping nucleus^60^, and later in peripheral tissues.

In teleosts, the pineal gland is considered the central pacemaker for daily rhythm^61^, though cyclic time-keeping gene expression is present in most tissues^62^; however, relatively little is known about the temporal pattern of expression of these genes in fish. Robust circadian expression patterns have been demonstrated of *per3* throughout the brain of zebrafish, with transcript expression peaking early in the light cycle, three hours after lights came on^63^. In nocturnal flatfish (*Solea senegalensis*), *per3* expression peaked at the transition between dark and light periods in the retina and optic tectum but showed no daily rhythm in the diencephalon or cerebellum^64^. One caveat in comparing these studies to the current investigation of midshipman is that they investigated entire brain regions such as the telencephalon, diencephalon, and rostral rhombencephalon rather than more focal sites comparable to the POA-AH and VMN. Still, as *per3* expression increases throughout the dark period, peaking during early light in these studies, our results are consistent with humming male VMN clocks being set to later subjective time than those in non-humming males.

As with *per3*, little is known about the temporal pattern of *Clock* expression in teleost brain. *Clock* transcript expression shows daily rhythms in zebrafish, with peaks in the eye, pineal gland, and whole brain near the onset of the dark period^65^. Similarly, in Nile tilapia (*Oreochromis niloticus*) expression of *clock1* transcripts in both hypothalamus and optic tectum peaked near the end of the light period^66^. As we found higher *Clock* expression in the POA-AH of humming males, our results could be consistent with these animals having either a later subjective clock, if non-humming males had not yet reached peak expression, or an earlier subjective clock if the non-humming fish had already reached peak expression and returned to baseline. In either scenario, our results show that humming males differ in circadian clock gene expression in both neuroendocrine and motor regions related to vocalization. This suggests that differences in internal subjective clock mechanisms may play an important role in determining if or when a fish will hum.

### 4.3 Conclusions

This study identifies several genes that are differentially expressed across fish that are actively vocalizing and those that are not, in both motor and neuroendocrine regions of the brain. Many of these genes, particularly in the VMN, fall under categories previously identified to be important for maintaining the precision and energetic demands of the extreme midshipman courtship hum, including synaptic transmission, ion channels and transport, hormone signaling, and metabolism and antioxidant activity. By directly comparing humming and non-humming animals, it was possible to expand on the list of candidate genes identified by previous studies that investigated gene expression differences across timescales relevant to humming behavior, but did not compare behaving animals, or that focused on reproductive behaviors that include humming, but did not test the effects of humming specifically. In addition to these categories, two genes involved in the molecular circadian clock were upregulated in humming males: *per3* and *Clock* in the VMN and POA-AH respectively. This result underscores the importance of circadian time keeping in regulating midshipman vocal behavior, a theme identified in prior studies.

## Acknowledgements

We would like to thank David Rose, Margaret Marchaterre, and Rich Moore for logistical assistance, Andrea Makowski for assistance in annotation, Joe Sisneros for assistance obtaining collection permits, and Eric Schuppe for comments on earlier versions of the manuscript. We also thank Cornell Computational Biology Service Unit for advice on data analysis. This work was supported by the National Science Foundation IOS-1457108 and IOS-1656664 (A.H.B.) and Cornell University Department of Neurobiology and Behavior (J.A.T.). The authors declare no conflict of interest.

## Data Availability Statement

The reads from each sample supporting the results of this article are available in the NCBI Sequence Read Archive database under BioProject accession number PRJNA665270.

